# Cadm1 regulates airway stem cell growth and differentiation via modulation of Stat3 activity

**DOI:** 10.1101/090761

**Authors:** Pooja Seedhar, Elizabeth K Sage, Sabari Vallath, Gabrielle Sturges, Adam Giangreco

## Abstract

Airway homeostasis, repair, and regeneration are imperfectly understood processes involving the proliferation and differentiation of endogenous lung stem cells. Here, we establish that epithelial Cell adhesion molecule 1 (Cadm1) regulates the growth and differentiation of airway basal cells, previously identified as lung stem cells. Immunohistochemistry and gene expression analysis reveals that Cadm1 is broadly expressed throughout the murine tracheobronchial epithelium, exhibits transient downregulation concomitant with airway injury, and is subsequently restored during basal cell differentiation. Using Cadm1 null (KO) and keratin 14 (K14)-specific Cadm1 overexpressing transgenic mice, we demonstrate that maintaining Cadm1 expression reduces basal stem cell proliferation after tracheal polidocanol injury, whereas Cadm1 deletion causes increased ciliated cell differentiation and sustained downstream Stat3 signalling. Altogether, this study defines a previously uncharacterised role for Cadm1 in directing airway basal cell homeostasis and repair via modulation of Stat3 activity.

## Introduction

Conducting airway injury due to infection, inflammation, toxicant exposure, and trauma causes disrupted mucociliary clearance and reduced barrier function that requires rapid repair and regeneration to restore lung health ^1-3^. Inefficient airway repair alters epithelial morphology and increases the risk of chronic lung diseases including bronchitis, chronic obstructive pulmonary disease (COPD), and cancer ^1,4-6^. Aberrant airway repair and inappropriate epithelial differentiation are also associated with an increased risk of lung tumorigenesis^5^.

Airway repair and regeneration are governed by endogenous lung stem cells capable of long term self-renewal and multipotent differentiation^7^. In human airways and the murine trachea these cells are defined as keratin 5 and keratin 14-expressing basal cells. Airway basal stem cell activation during epithelial repair and regeneration involves upregulation of the Wnt/beta-catenin, Notch, and IL6/Stat3 signalling pathways ^8-10^. ^5,11-14^. In addition, responding appropriately to local cell-cell and cell-matrix interactions is an essential component of airway basal stem cell function ^15^. Despite this, the mechanisms by which local cell adhesion molecules direct basal cell phenotypes remain largely unknown.

In previous studies, we and others have shown that epithelial Cell adhesion molecule 1 (Cadm1) expression regulates epithelial cell growth in a variety of tissues ^16-24^. Cadm1 is a nectin-like adhesion molecule responsible for maintaining calcium-independent cell-cell adhesions ^21^. Cadm1 mediates homotypic and heterotypic intercellular interactions via nectin 3 and Necl1, has been shown to enhance E-cadherin dependent adherens junction stability, and indirectly modulates cytoskeletal RhoGTPase activity via cytoplasmic PDZ domain interactions ^21,22,24^. Recently, we also established that Cadm1 inhibits human squamous cell carcinoma growth and metastases by reducing downstream Stat3 activity ^17^.

Here, we investigate the role of Cadm1 in regulating airway basal stem cell growth and differentiation in the context of lung homeostasis and repair. We establish that Cadm1 is broadly expressed throughout murine airways and exhibits rapid downregulation upon epithelial injury. Our results indicate that transient Cadm1 loss is a necessary component of normal airway repair, whilst Cadm1 restoration modulates ciliated cell differentiation and downstream Stat3 signalling.

## Materials and Methods

### In vivo procedures

Adult (2-6 month old) K14-Cadm1 transgenic (TG, strain 7248A) and Cadm1-null (KO) mice were generated as previously described,18,24, housed in individually ventilated cages on a 12 hour light-dark cycle and allowed access to food and water ad libitum. Wild type (WT) littermates were used as controls. Genotyping was achieved by PCR amplification of ear biopsy DNA as previously described^24^. For airway regeneration studies, mice were anaesthetized with isoflurane and their tracheas damaged by oropharyngeal instillation of 15µl 2% polidocanol (Sigma, UK). All mice additionally received a single intraperitoneal injection of BrdU (10mg/kg body weight; Invitrogen UK) two (transgenic) or six hours prior to sacrifice by sodium pentobarbital overdose. All experiments were performed under the terms of a UK Home Office license.

### Tissue preparation and antibody staining

Tracheal tissue sections were fixed overnight at 4 neutral buffered formalin, processed, sectioned and stained with haematoxylin and eosin or primary antibodies using standard conditions^5^. Tracheal whole mounts were fixed, microdissected to expose the luminal airway surface, and stained using lung whole mount conditions^9^. Primary antibodies included keratin 14 (rabbit, Covance), keratin 5 (rabbit, Covance), BrdU (rat, Abcam), Cadm1 (chicken, MBL), FoxJ1 (mouse, eBioscience), acetylated tubulin (ACT, mouse, Sigma), Club cell secretory protein (CCSP, goat, gift of B Stripp), and phospho-Stat3 (pStat3, rabbit, Cell Signalling). Species-appropriate secondary antibodies were directly conjugated to either Alexfluor 488 or 555 dyes (all 1:300; Invitrogen, UK).

### Imaging

A Nanozoom system was used to capture images of H+E stained slides (Hammatsu, Japan). A Zeiss Axioscope 2 with QICAM camera or Leica TCS SP8 confocal were used for fluorescent imaging. Tracheal whole mount images were obtained using a Zeiss SterEO Lumar V.12 automated imaging system. Images were processed using Zeiss Axiovision, Leica LAS-AF, FIJI, and Adobe Photoshop software (to rotate, crop, and adjust brightness and contrast).

### Morphometry and statistical analysis

We used Volocity and FIJI image analysis software to quantify BrdU abundance and K14-expressing cell coverage as a function of epithelial surface area in tracheal wholemounts. We also used Volocity to assess airway epithelial height in haematoxylin and eosin (H+E) stained tissue sections.

Pairwise *t*-tests were used for all statistical comparisons. Statistical significance was accepted at *p* < 0.05 for all analyses. All statistical analyses were performed using GraphPad Prism and Microsoft Excel.

## Results

### Tracheal injury is associated with reduced Cadm1 expression

In the steady state tracheal epithelium, Cadm1 was readily detected in both basal stem cells (GFP+/ lectin+) and non-basal cell types (GFP-/lectin+, GFP-/lectin-) by gene expression profiling (Figure 1A, GSE15724,^8^). Separate gene expression microarray analyses revealed that Cadm1 levels were significantly reduced 48 hours after intratracheal SO2 injury (Figure 1B, GSE69056, ^11^) and 72 hours after oropharyngeal polidocanol-mediated tracheal damage (Figure 1C, GSE17268, ^25^). Both polidocanol and SO2 injury caused rapid epithelial injury and cell desquamation, activation of endogenous basal stem cell proliferation within 72 hours, and restoration of a differentiating epithelium within 2 weeks. Whole trachea quantitative PCR analysis confirmed that Cadm1 gene expression was significantly reduced 3 days after polidocanol administration and restored within 2 weeks post-injury (Supplemental Figure 1).

**Figure 1:**
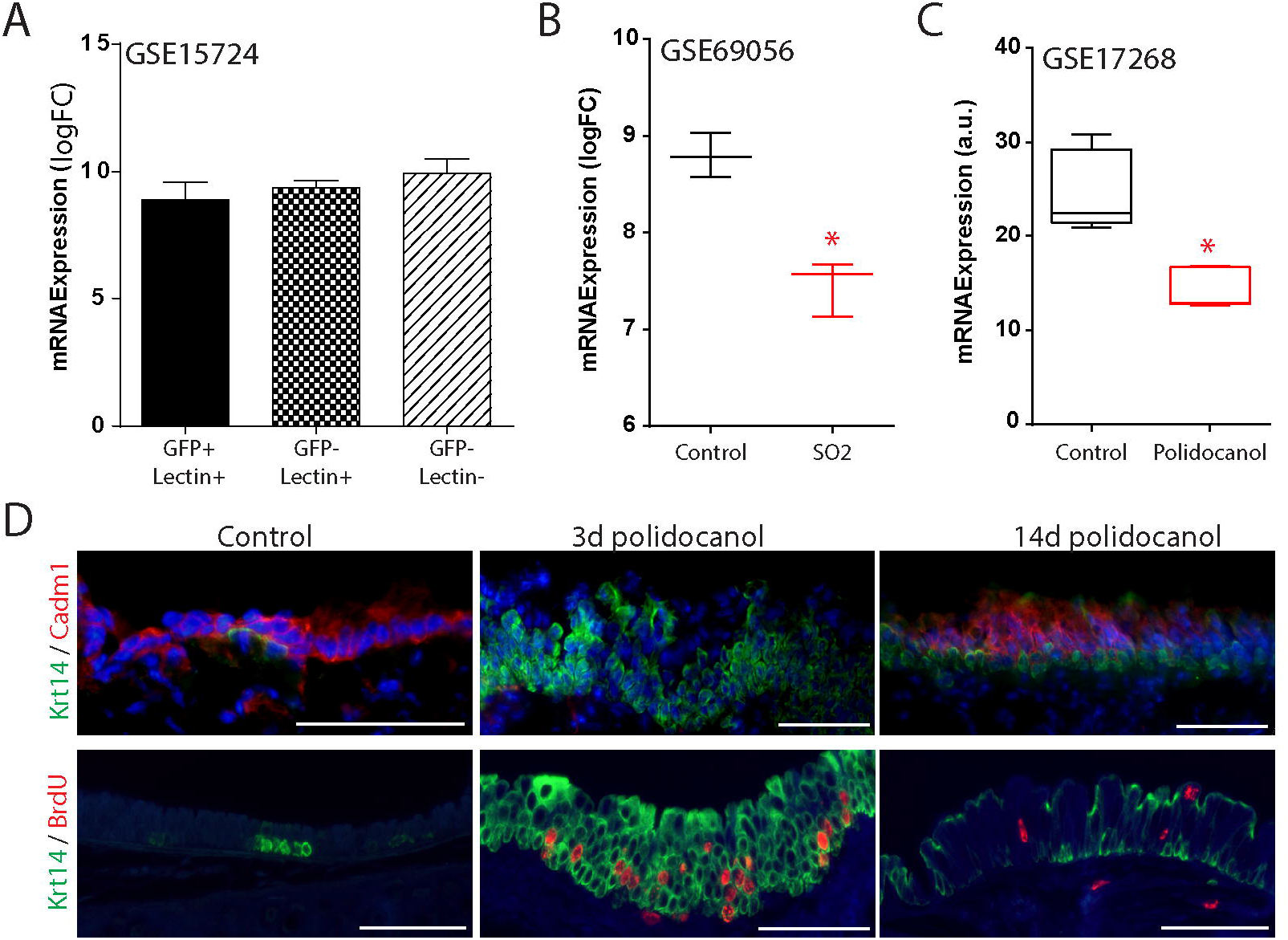
Cadm1 expression is reduced during tracheal injury and repair. (A-C) Relative Cadm1 gene expression in microarray datasets of GFP (+) and/or lectin (+) lung cells in the absence of injury (GSE15724, A), sulphur dioxide (SO2) treated versus uninjured whole tracheas (GSE69056, B), and polidocanol treated versus uninjured tracheas (GSE17268, C). (D) Immunostaining identifies Cadm1 (red), BrdU (red), and keratin 14 (K14, green) positive cells in normal and regenerating airways 3 and 14 days after polidocanol injury. Scale bars in (D) are 100µm * denotes p<0.05 (B, C).

To investigate differences in Cadm1 expression during airway injury, we used immunohistochemistry to assess the abundance and distribution of Cadm1 before and after polidocanol delivery (Figure 1D). Consistent with microarray data in Figure 1A, in the absence of injury Cadm1 was readily observed throughout proximal and bronchiolar airways in all keratin 14-expressing (K14 (+)) basal progenitor cells, secretory and ciliated cells. Three days after polidocanol administration, Cadm1 exhibited robust downregulation concomitant with extensive K14 (+) basal stem cell hyperplasia and proliferation. Fourteen days after polidocanol injury, Cadm1 was again detected in K14-negative and K14 (+) basal stem cells. This restoration of Cadm1 immunoreactivity was also associated with reduced BrdU (+) cell abundance and K14 (+) cell hyperplasia (Figure 1D).

### Sustained Cadm1 expression disrupts basal stem cell-mediated airway repair

To determine whether Cadm1 loss was necessary for tracheal repair, K14-Cadm1 transgenic (TG) mice were treated with oropharyngeal polidocanol. In the absence of polidocanol injury, there were no differences in cellularity, differentiation, or proliferation between WT and TG airways (Figure 2A, B). After polidocanol injury, histological examination revealed that TG airways were comprised of large regions of poorly regenerated, squamous epithelium when compared with WT controls (Figure 2A). This finding was corroborated by differences in airway epithelial height observed when comparing polidocanol treated TG mice with their WT littermates (Supplemental Figure 2). Despite significant differences in repair 3 days after polidocanol administration, WT and TG tracheal epithelial histology was indistinguishable 14 days after polidocanol exposure (Figure 2A). Thus, sustained expression of Cadm1 in TG mice delays but does not completely block airway epithelial repair. These results are consistent with our previous observation regarding epidermal Cadm1 expression and skin wound healing ^24^.

**Figure 2:**
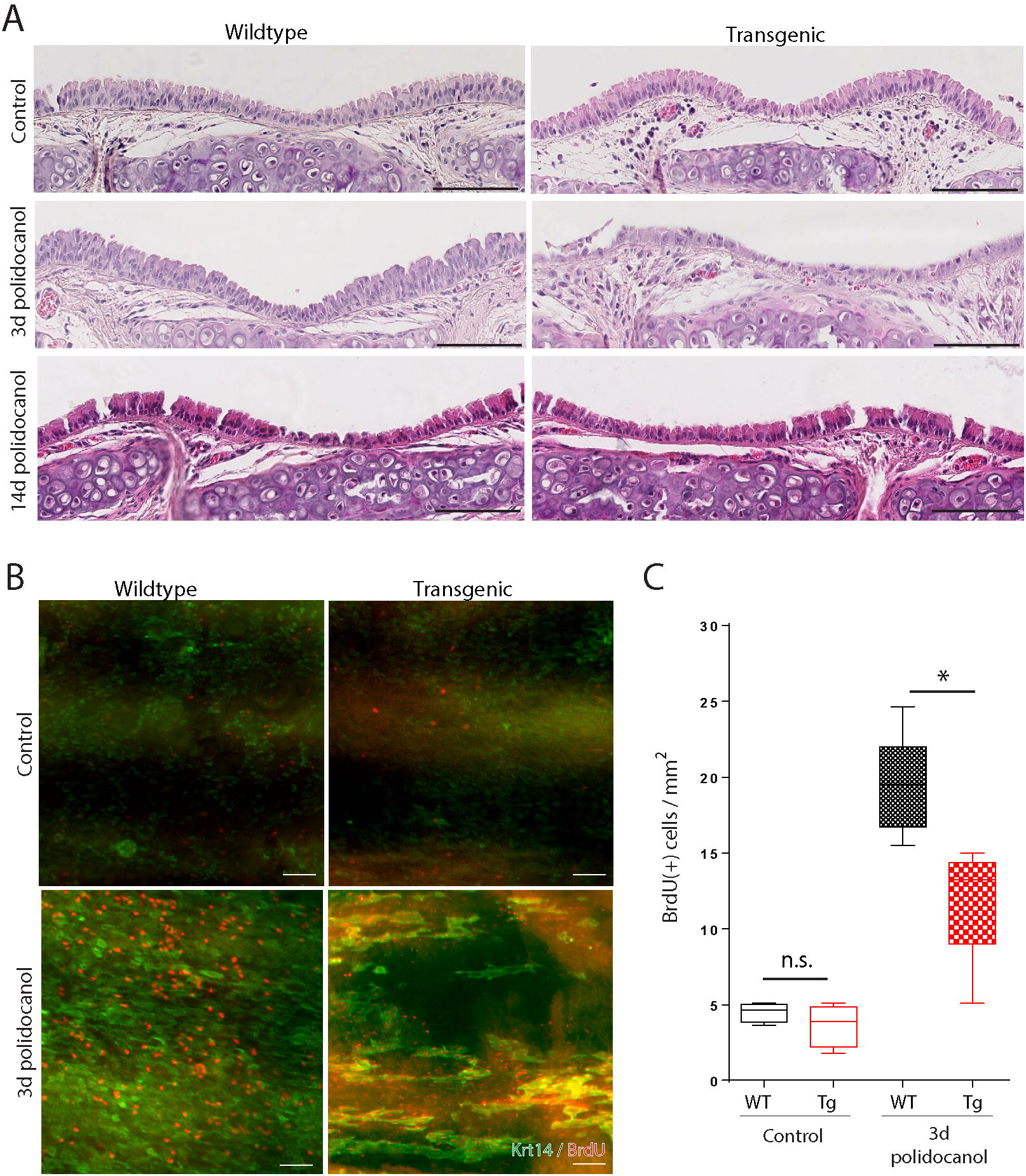
Sustained Cadm1 expression delays normal tracheal repair. (A) Representative H&E staining of uninjured (control) or regenerating wildtype and K14-Cadm1 transgenic airways 3 and 14 days after polidocanol injury. (B) Whole mount immunostaining for K14 (green) plus BrdU (red) in control or polidocanol treated wildtype and K14-Cadm1 airways. (C) Quantitation of BrdU (+) cell abundance in uninjured and polidocanol treated wildtype (WT, black) and K14-Cadm1 transgenic (TG, red) airway wholemounts. Scale bars (A, B) are 100µm; * denotes p<0.05 (C).

To examine the mechanism by which sustained Cadm1 expression affected airway stem cell-mediated repair, proliferating K14(+) basal cells were labelled with BrdU 3 days after injury (Figure 2B). Analysis of tracheal whole mount sections revealed significantly fewer BrdU(+) basal cells in repairing TG airways compared with WT littermates (19 BrdU(+) cells/mm2 in wild type versus 13 BrdU(+) cells/mm2 in transgenics; Figure 2B, C). Mitotic, hyperplastic K14 (+) cells were additionally observed in cohesive cell clusters, raising the possibility that different subpopulations of basal cells may respond differently to sustained Cadm1 expression^26^.

### Cadm1 loss promotes increased basal cell proliferation and ciliated cell differentiation

To establish whether Cadm1 loss was sufficient to promote airway progenitor cell hyperplasia we compared WT and Cadm1-null (KO) airways. In the absence of airway injury, there were no clear histological differences between WT and KO samples (Figure 3A). Epithelial height and K14 (+) cell hyperplasia were also similar in uninjured animals (Figure 2B, Supplemental Figure 3). However, we did observe increased BrdU (+) cell abundance in KO airways compared with WTs (Figure 3B, C). Thus, deletion of Cadm1 is sufficient to promote increased airway proliferation but not epithelial hyperplasia in the absence of injury. Three and 14 days after polidocanol injury, there were no differences in WT and KO epithelial height, K14(+) cell hyperplasia, or BrdU(+) cell abundance (Figure 3A-C and data not shown). These results were consistent with our finding that loss of endogenous Cadm1 expression accompanies normal airway injury and repair (Figure 1).

**Figure 3:**
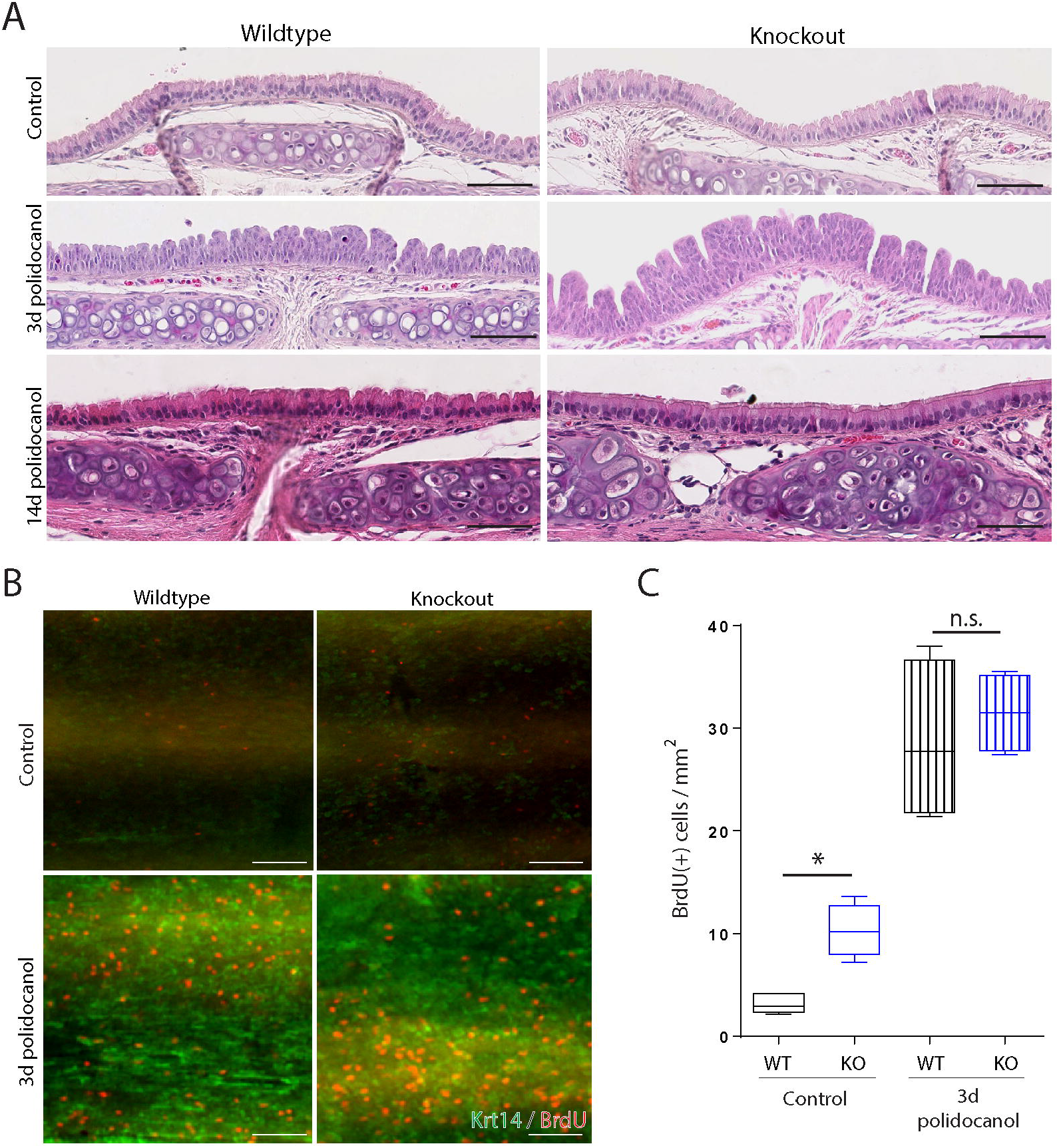
Cadm1 KO mice exhibit increased stem cell proliferation in the absence of injury. (A) Representative H&E staining of uninjured (control) or regenerating wildtype and Cadm1 knockout (KO) airways 3 and 14 days after polidocanol injury. (B) Whole mount immunostaining for K14 (green) plus BrdU (red) in control or polidocanol treated wildtype and Cadm1 KO airways. (C) Quantitation of BrdU (+) cell abundance in uninjured and polidocanol treated wildtype (WT, black) and Cadm1 KO (KO, blue) airway wholemounts. Scale bars (A, B) are 100µm; * denotes p<0.05 (C).

Unexpectedly, a histological examination of WT and KO tracheas 14 days post-polidocanol injury revealed significantly increased ciliated cell differentiation in Cadm1 KO airways (Figure 3A). We therefore assessed mucosecretory, basal, and ciliated cell abundance using dual immunofluorescent immunostaining for Club cell secretory protein (CCSP) plus acetylated tubulin (ACT), or keratin 5 plus FoxJ1 (Figure 4). Whilst both WT and KO airways were comprised of a mix of secretory, basal, and ciliated cells in the absence of injury, the percentage of FoxJ1(+) and/or ACT(+) cells in the airways of KO mice were significantly increased 14 days after polidocanol exposure (Figure 4A, B). On the other hand, CCSP (+) cells were virtually absent from KO airways following polidocanol injury and repair (Figure 4A). Thus, Cadm1 deletion regulates the differentiation of basal stem cells towards a ciliated cell fate after tracheal polidocanol injury.

**Figure 4:**
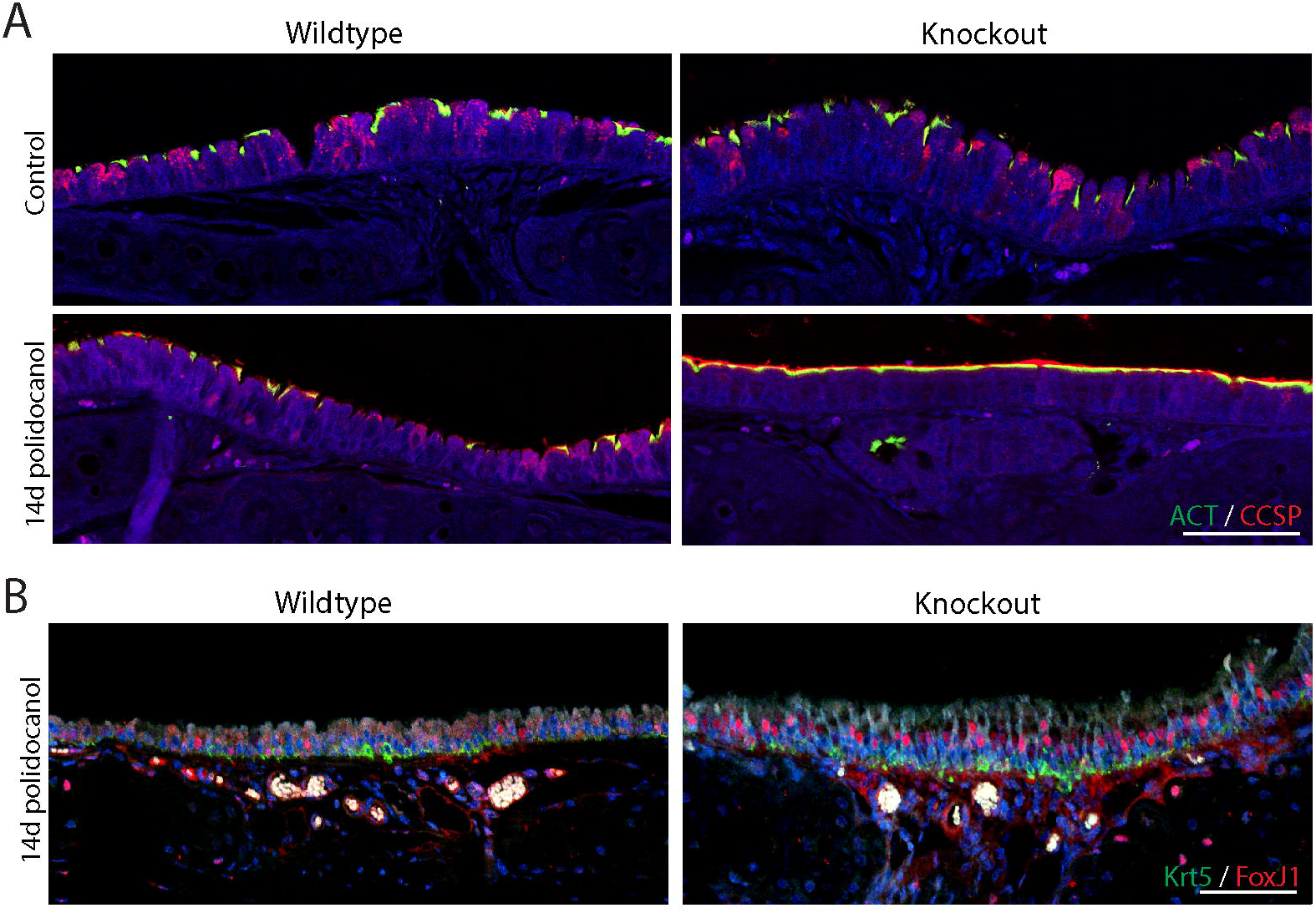
Cadm1 deletion promotes increased ciliated cell differentiation. (A) Immunostaining for secretory (CCSP, red) plus ciliated (ACT, green) cell abundance in uninjured control or polidocanol treated wildtype and Cadm1 knockout airways recovered for 14 days. (B) Immunostaining for basal (Krt5, green) plus ciliated FoxJ1-expressing (red) cell abundance in polidocanol injured wildtype and Cadm1 KO airways recovered for 14 days. Scale bars (A, B) are 100µm.

### Cadm1 regulates downstream Stat3 activity

Given previous reports that sustained Stat3 activity can promote regeneration of airway ciliated cells at the expense of secretory cell lineages ^11^, is required for bronchiolar epithelial repair^27^, and is regulated by Cadm1 ^17^, we investigated whether Stat3 activity was altered in Cadm1 TG and KO airways before and after polidocanol injury. Using immunohistochemistry, we observed abundant pStat3 immunostaining in K5 (+) basal cells of KO animals before, during, and 14 days after polidocanol exposure (Figure 5). In contrast, pStat3 immunostaining was only present in WT airway basal cells 3 days after polidocanol injury (Figure 5). Consistent with this finding, Cadm1 TG airways had very few pStat3 (+) cells before, during, or after polidocanol injury and repair (Supplemental Figure 4). Thus, Cadm1 deletion promotes sustained downstream Stat3 activation, whilst Cadm1 overexpression inhibits Stat3 signalling. Together, these results provide a plausible mechanism by which the intercellular adhesion molecule Cadm1 regulates both basal stem cell proliferation and ciliated cell differentiation via modulation of downstream Stat3 activity.

**Figure 5:**
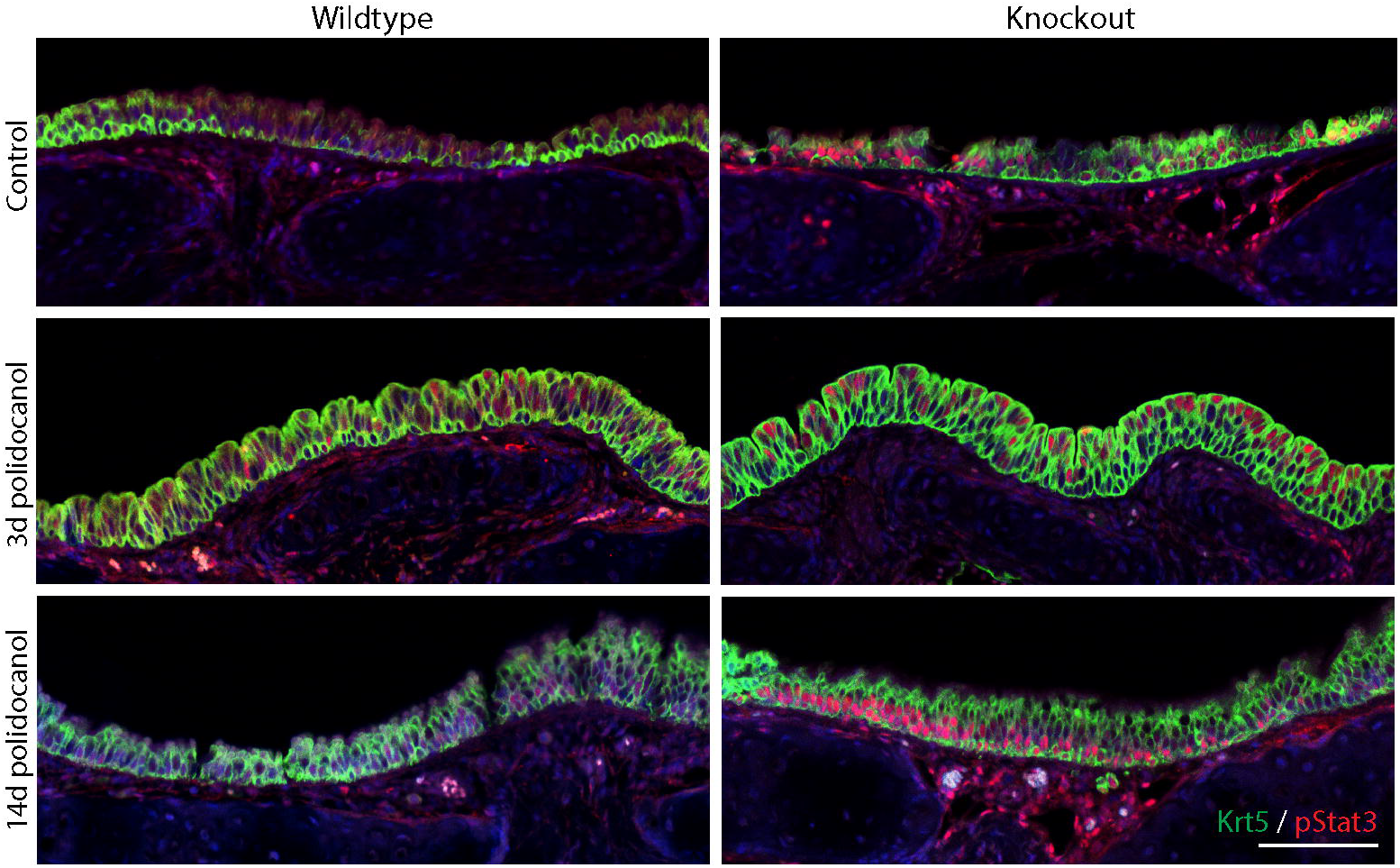
Cadm1 KO airways exhibit sustained Stat3 activity. Phospho-Stat3 (pStat3, red) plus keratin 5 (Krt5, green) staining in uninjured and polidocanol treated wildtype and Cadm1 KO airways recovered for 3 or 14 days. Scale bar is 100µm.

## Discussion

This study defines the epithelial cell adhesion molecule Cadm1 as an important regulator of airway homeostasis and repair. In the absence of injury, Cadm1 is abundantly expressed throughout conducting airways whilst transient loss of Cadm1 accompanies tracheal damage and basal stem cell-mediated repair. Using genetically modified mouse models, we establish that Cadm1 downregulation is necessary for stem cell proliferation and that sustained Cadm1 loss promotes increased ciliated cell differentiation. Finally, we provide evidence that Cadm1 regulates airway stem cell behaviour via modulation of downstream Stat3 activity.

Our data are consistent with identified roles for Cadm1 in other tissues and for Stat3 in lung repair and differentiation. In other tissues, Cadm1 suppresses cell scattering^22^, promotes epithelial cell tubulogenesis^23^, and directs stem cell mediated wound healing^24^. Likewise, Stat3 signalling regulates epithelial cell migration, is required for bronchiolar epithelial repair, and promotes regeneration of ciliated cells from basal stem cells,11,27. Given our recently established functional links between Cadm1 and Stat3 in squamous cell carcinoma^17^, it is likely that Cadm1 functions in airways to indirectly modulate IL6-dependent Stat3 activation. Under homeostatic conditions, Cadm1 is likely to suppress Stat3 activation, thereby restricting airway stem cell turnover. Airway injury, resulting in Cadm1 loss as well as local IL-6 secretion, would promote de-repression of Stat3 activation and increased cell proliferation and ciliated cell differentiation^11^.

Overall, this study highlights the importance of Cadm1 in airway repair and regeneration. Although focussed on a single cell adhesion molecule, these results are consistent with earlier reports on the role of cadherins, integrins, and other junctional adhesion molecules in the homeostasis, repair and regeneration of airways,1,28. Similarly, cell adhesion molecules are known to play major roles in disease pathogenesis by influencing intrinsic cellular signalling and the interaction of progenitor cells with their surrounding environment^29^. Thus, our results highlight a plausible mechanism by which aberrant basal stem cell activation during injury and repair could contribute to chronic lung pathologies such as cancer.

## Acknowledgements

We acknowledge the University College London biological services unit for animal care and husbandry and members of the UCL Lungs for Living Research Centre who read the manuscript. AG is the recipient of a European Research Council Starting Investigator award and EKS is the recipient of a Medical Research Council Clinical Training Fellowship and NIHR clinical lectureship.

### Author contributions

PS designed, performed and analysed experiments; EKS, SV, and GS performed and analysed experiments; AG conceived, designed, performed and analysed experiments and wrote the paper.

### Conflict of interest

None of the authors declare any conflicts of interest.

## References

1 Crosby, L. M. & Waters, C. M. Epithelial repair mechanisms in the lung. Am J Physiol Lung Cell Mol Physiol 298, L715–731, doi:10.1152/ajplung.00361.2009 (2010).

2 Rackley, C. R. & Stripp, B. R. Building and maintaining the epithelium of the lung. J Clin Invest 122, 2724–2730, doi:10.1172/JCI60519 (2012).

3 Bertoncello, I. & McQualter, J. L. Lung stem cells: do they exist? Respirology 18, 587–595, doi:10.1111/resp.12073 (2013).

4 Puddicombe, S. M. et al. Involvement of the epidermal growth factor receptor in epithelial repair in asthma. Faseb J 14, 1362–1374 (2000).

5 Giangreco, A. et al. beta-Catenin determines upper airway progenitor cell fate and preinvasive squamous lung cancer progression by modulating epithelial-mesenchymal transition. J Pathol 226, 575–587, doi:10.1002/path.3962 (2012).

6 Fischer, B. M. et al. ErbB2 activity is required for airway epithelial repair following neutrophil elastase exposure. Faseb J 19, 1374–1376, doi:10.1096/fj.04-2675fje (2005).

7 Hogan, B. L. et al. Repair and regeneration of the respiratory system: complexity, plasticity, and mechanisms of lung stem cell function. Cell Stem Cell 15, 123–138, doi:10.1016/j.stem.2014.07.012 (2014).

8 Rock, J. R. et al. Basal cells as stem cells of the mouse trachea and human airway epithelium. Proc Natl Acad Sci U S A 106, 12771–12775, doi:10.1073/pnas.0906850106 (2009).

9 Giangreco, A. et al. Stem cells are dispensable for lung homeostasis but restore airways after injury. Proc Natl Acad Sci U S A 106, 9286–9291, doi:10.1073/pnas.0900668106 (2009).

10 Hong, K. U., Reynolds, S. D., Watkins, S., Fuchs, E. & Stripp, B. R. Basal cells are a multipotent progenitor capable of renewing the bronchial epithelium. Am J Pathol 164, 577–588, doi:10.1016/S0002-9440(10)63147-1 (2004).

11 Tadokoro, T. et al. IL-6/STAT3 promotes regeneration of airway ciliated cells from basal stem cells. Proc Natl Acad Sci U S A 111, E3641–3649, doi:10.1073/pnas.1409781111 (2014).

12 Rock, J. R. et al. Notch-dependent differentiation of adult airway basal stem cells. Cell Stem Cell 8, 639–648, doi:10.1016/j.stem.2011.04.003 (2011).

13 Tsao, P. N. et al. Notch signaling controls the balance of ciliated and secretory cell fates in developing airways. Development 136, 2297–2307, doi:10.1242/dev.034884 (2009).

14 Vermeer, P. D. et al. Segregation of receptor and ligand regulates activation of epithelial growth factor receptor. Nature 422, 322–326, doi:10.1038/nature01440 (2003).

15 Rock, J. R., Randell, S. H. & Hogan, B. L. Airway basal stem cells: a perspective on their roles in epithelial homeostasis and remodeling. Dis Model Mech 3, 545–556, doi:10.1242/dmm.006031 (2010).

16 Murakami, Y. Involvement of a cell adhesion molecule, TSLC1/IGSF4, in human oncogenesis. Cancer Sci 96, 543–552, doi:10.1111/j.1349-7006.2005.00089.x (2005).

17 Vallath, S. et al. CADM1 inhibits squamous cell carcinoma progression by reducing STAT3 activity. Sci Rep 6, 24006, doi:10.1038/srep24006 (2016).

18 Fujita, E. et al. Oligo-astheno-teratozoospermia in mice lacking RA175/TSLC1/SynCAM/IGSF4A, a cell adhesion molecule in the immunoglobulin superfamily. Mol Cell Biol 26, 718–726, doi:10.1128/MCB.26.2.718-726.2006 (2006).

19 van der Weyden, L. et al. Loss of TSLC1 causes male infertility due to a defect at the spermatid stage of spermatogenesis. Mol Cell Biol 26, 3595–3609, doi:10.1128/MCB.26.9.3595-3609.2006 (2006).

20 Rikitake, Y., Mandai, K. & Takai, Y. The role of nectins in different types of cell-cell adhesion. J Cell Sci 125, 3713–3722, doi:10.1242/jcs.099572 (2012).

21 Takai, Y., Miyoshi, J., Ikeda, W. & Ogita, H. Nectins and nectin-like molecules: roles in contact inhibition of cell movement and proliferation. Nat Rev Mol Cell Biol 9, 603–615, doi:10.1038/nrm2457 (2008).

22 Masuda, M. et al. Tumor suppressor in lung cancer (TSLC)1 suppresses epithelial cell scattering and tubulogenesis. J Biol Chem 280, 42164–42171, doi:10.1074/jbc.M507136200 (2005)

23 Ito, A. et al. SgIGSF is a novel biliary-epithelial cell adhesion molecule mediating duct/ductule development. Hepatology 45, 684–694, doi:10.1002/hep.21501 (2007)

24 Giangreco, A., Jensen, K. B., Takai, Y., Miyoshi, J. & Watt, F. M. Necl2 regulates epidermal adhesion and wound repair. Development 136, 3505–3514, doi:10.1242/dev.038232 (2009).

25 Giangreco, A. et al. Myd88 deficiency influences murine tracheal epithelial metaplasia and submucosal gland abundance. J Pathol 224, 190–202, doi:10.1002/path.2876 (2011).

26 Watson, J. K. et al. Clonal Dynamics Reveal Two Distinct Populations of Basal Cells in Slow-Turnover Airway Epithelium. Cell Rep 12, 90–101, doi:10.1016/j.celrep.2015.06.011 (2015).

27 Kida, H. et al. GP130-STAT3 regulates epithelial cell migration and is required for repair of the bronchiolar epithelium. Am J Pathol 172, 1542–1554, doi:10.2353/ajpath.2008.071052 (2008).

28 Nawijn, M. C., Hackett, T. L., Postma, D. S., van Oosterhout, A. J. & Heijink, I. H. E-cadherin: gatekeeper of airway mucosa and allergic sensitization. Trends Immunol 32, 248–255, doi:10.1016/j.it.2011.03.004 (2011).

29 Watt, F. M. Role of integrins in regulating epidermal adhesion, growth and differentiation. Embo J 21, 3919–3926, doi:10.1093/emboj/cdf399 (2002).

